# Chromatic contrasts predict human target detection in a natural search task

**DOI:** 10.64898/2026.07.11.737926

**Authors:** Thomas E. White, Bibiana Rojas, Johanna Mappes, Darrell J. Kemp

## Abstract

Natural scenes are visually complex, with dynamic variation in colour and luminance presenting challenges for detecting salient objects. However, the cues guiding human detection under such natural conditions remain largely untested. We placed custom-built coloured targets in a forest-like setting and quantified their contrasts against natural backgrounds using human visual models. Across human observers (n = 51), detection performance rose sharply with hue and saturation contrast, whereas luminance contrast—long considered a primary driver of visual search—contributed little explanatory power. Strikingly, even large changes in ambient light chromaticity barely altered detection, despite the patchy illumination of the habitat. Under these naturalistic conditions, hue and saturation contrasts were stronger predictors of detection than luminance contrast, providing a rare, field-based test of visual models widely applied across human and animal vision research.

## Introduction

The world is a visually noisy place. Natural scenes are alive with shimmering fields of colour and luminance in motion (Geisler, 2008). This vibrant chaos presents a fundamental challenge for visual systems: the extraction of salient cues to guide higher-level tasks such as foraging, mate selection, and predator avoidance (Forster, 2023; Kemp, 2007; Rojas et al., 2014). In humans, the earliest stages of perception are well characterised. Our three differentially-sensitive cone receptors are distributed unevenly across the retina, and are processed in two opponent chromatic (‘colour’) channels and one non-opponent luminance (‘brightness’) channel (Gegenfurtner & Kiper, 2003). This affords access to the basic cues of hue, chroma, and luminance which are, in effect, the raw sense data that ultimately guide perception. The value assigned to these cues is not, however, equally apportioned across tasks. The relative stability of colour information, for example, sees it drawn on to inform object identification and categorisation, visual search, and memory (Kelber & Osorio, 2010). Sensing motion, texture, and depth, by contrast, are chiefly luminance-guided, owing to the heightened speed and sensitivity of this channel (Delorme et al., 2000). This initial mapping of cues to tasks is common to most visual animals, but its operation and effectiveness in the chaos of the natural world is much less well understood (Osorio & Vorobyev, 2005).

Human colour vision is uniquely well characterised among animals, such that it can be reliably (albeit imperfectly) represented in human-centred visual models (Ibraheem et al., 2012). Exemplars include CIELab and its derivatives, which see widespread use in visual design and display calibration. The ultimate objective of such models is to faithfully represent the range of colours perceivable by viewers, and so translate a nuanced understanding of visual physiology into practical tools that predict our perception of and interaction with stimuli and environments (Kemp et al., 2015). Despite considerable progress in linking physiology to sensory experience via such models, efforts to validate their predictions through tests of visual performance ‘in the wild’ are few. Indeed, the near sum-total of knowledge to date is laboratory based, under artificial conditions (Ibraheem et al., 2012; Kemp et al., 2015; Renoult et al., 2015). Such work, while invaluable, has positioned the broader ecology of human colour vision as fertile ground in need of study.

The consequences of this narrow view stretch well beyond human domains. Human perception is uniquely tractable, and the fundamentals of colour sensation are shared across most animals (Kelber & Osorio, 2010; Osorio & Vorobyev, 2005). This means that insights garnered from the human experience are often extrapolated to inform non-human visual systems and models, to great effect. Psychophysical tests of human colour vision are routinely adapted for use in non-human primates (Kawamura, 2016), while comparative work continues to probe the unique (among mammals) evolutionary origins of primate trichromacy (Surridge et al., 2003). Further afield, studies on animal signalling and camouflage regularly use human viewers as proxies for mammalian and avian predators (Xiao & Cuthill, 2016). This breadth of work argues for a twofold value of tests of human visual performance in the wild. One is the direct insight into the adaptive value and ecology of human colour vision, extending to contemporary technological challenges in signalling and display design (Ibraheem et al., 2012; Kawamura, 2016). The other is the centrality of human colour vision to taxonomically and conceptually broader questions in physiology, ecology, and evolution (Endler & Mappes, 2017; Kemp et al., 2015).

Here we set out to test the visual cues that govern human detection of objects in natural, forest-like environments. Building on White et al. (2017), but using targets specifically designed to vary across a broader range of visual contrasts, we tested how well differences in these first-order visual features predict detection performance under field conditions. Based on theory and earlier laboratory work, we predicted that chromatic cues—hue and saturation—would be stronger predictors of detection than luminance because colour information tends to remain more stable than luminance under variable natural illumination (Gegenfurtner & Kiper, 2003; Kelber & Osorio, 2010; Osorio & Vorobyev, 2005). By linking visual-model predictions to realised performance in a natural environment, our study provides a rare field-based assessment of the visual cues associated with human target detection.

## Methods

### Visual stimulus design

We designed and constructed small visual targets with the intent of creating stimuli which spanned a controlled but broad range of both chromatic (hue and saturation) and achromatic (lightness) variation. To achieve this, we used short lengths of wooden dowel (15 mm diameter, 30 mm length) as a base, and wrapped each first in white filter paper and then in either a single or double-layer of Rosco e-colour+ filters cut to size depending on their treatment. Our seven stimuli were comprised of three ‘light’ coloured targets —yellow, green, and blue — built of a single layer of filter sheet; three ‘dark’ coloured targets using the same coloured filters though applied in a double layer; and one ‘control’ target designed to match the average visual background, built of a double-layer of dark brown filter sheet (Fig. 1a-b).

**Figure 1:**
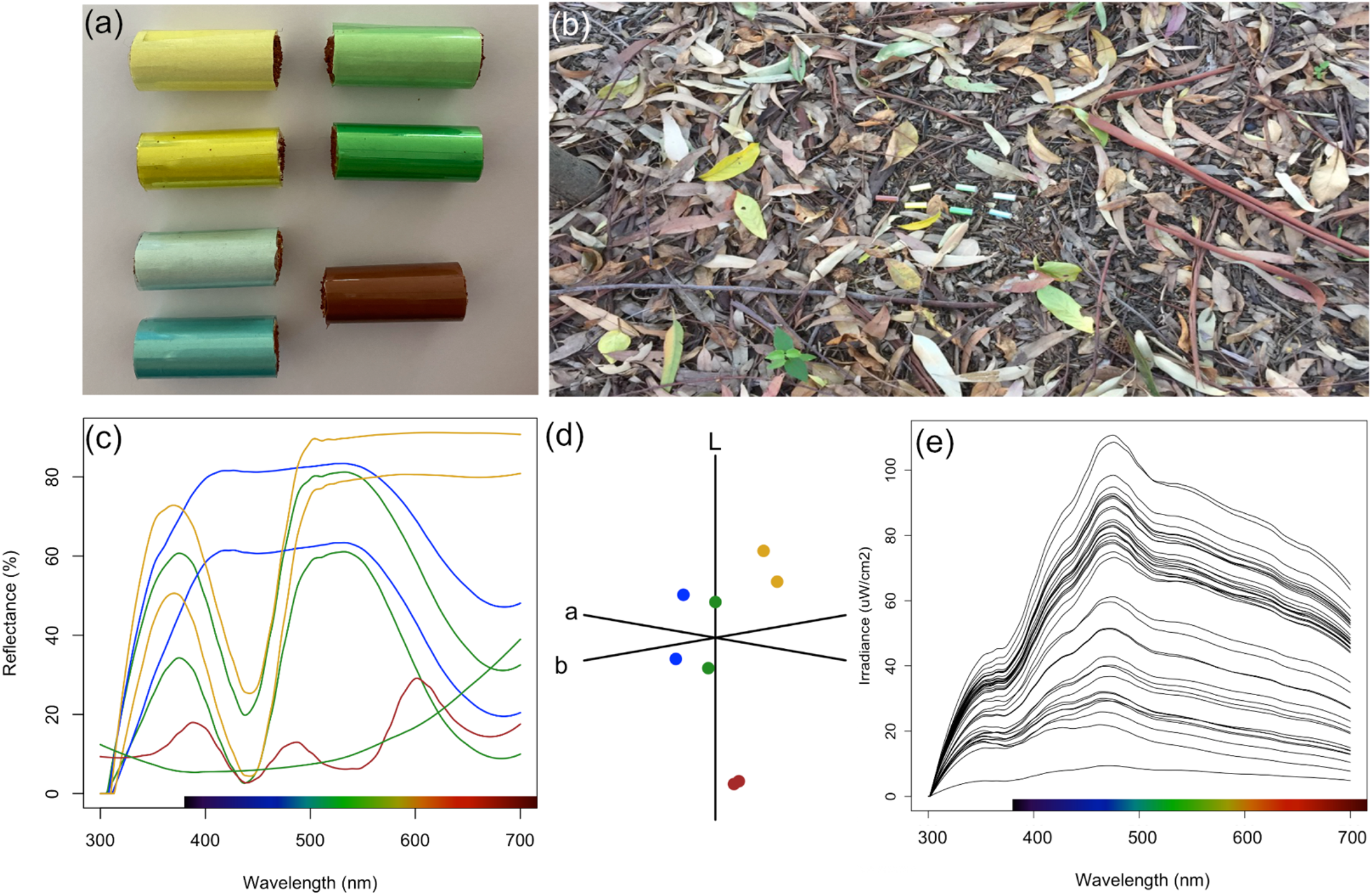
Visual properties of the experimental stimuli and light environment. (a) Photographs of the seven experimental targets used in detection assays: three chromatically matched hues (yellow, green, blue), each presented as a “light” (single-layer) and “dark” (double-layer) variant, plus a background-matching brown control. (b) Example of targets deployed on the natural search substrate during a field trial, illustrating typical visual clutter and patchy illumination. (c) Mean reflectance spectra for the seven target types, showing chromatic matching within hues and the imposed lightness contrast between “light” and “dark” variants. (d) Targets plotted in CIELCh colour space (D65 illuminant; 10° standard observer), illustrating relative differences in hue, chroma, and lightness. (e) Ambient irradiance spectra recorded during field assays, showing variation in the forest-like light environment under which participants searched for targets.

To estimate the human-perceived visual contrast achieved by our target set we used a combination of spectrophotometry and visual modelling as per White et al. (2017). Briefly, we used an OceanOptics USB4000 spectrometer coupled with a PX-2 pulsed xenon light source and 500 µm optical probes (set at 90° and 45° for light and collector, respectively) to record the reflectance spectrum of three replicate targets of each type, before averaging the resulting spectra (Fig. 1c). For each measurement we used a 99% diffuse ‘spectralon’ standard (Labsphere, New Hampshire) as a white reference, blocked the collector port to set a dark reference, and we specified a boxcar width of 10 and integration time of 200 ms per scan. We subsequently binned all spectra at 1 nm wavelength intervals and used minor LOESS (alpha = 0.15) to remove noise. In addition to our targets, we collected a sample of background material (hard tree-bark mulch, described below), and collected 10 spectra which we averaged to generate a representative background spectrum. Because we characterised the background with a single averaged spectrum, our contrasts describe differences from the mean background rather than the full heterogeneity of the natural substrate, such that targets matching the average background may still differ from some local patches (and vice versa). We then used the standard calculations for CIELab model and its CIELCh cylindrical transformation to estimate the subjective contrasts presented by our targets in the environment (Westland et al., 2012). We used the 10-degree standard observer colour-matching function under a D65 illuminant, and we calculated contrasts in each of hue, saturation, and lightness as the simple difference between targets and backgrounds in each dimension (Westland et al., 2012; White et al., 2017). Since our stimuli subtended less than 1° of visual angle, our use of the standard 10° observer may introduce some error in absolute colour estimates. Its impacts are negligible, however, as all stimuli were analysed with the same parameterisation and our conclusions depend on relative differences among treatments rather than absolute perceptual coordinates. For the same reason, our assumptions of a D65 illuminant and of predominantly diffuse reflectance—despite minor specular components on some targets—affect absolute rather than relative estimates. Throughout, we use ‘contrast’ to denote the difference between target and background along a given CIELAB dimension, and we treat the achromatic axis (CIELAB L*) as lightness, using ‘luminance contrast’ for contrast along this achromatic dimension. We ran all spectral processing and visual modelling in R (v. 4.3.1) using the packages ‘lightr’ (1.7.1) and ‘pavo’ (v 2.10.0) (Gruson et al., 2019; Maia et al., 2019, p. 2; Maia & White, 2018; R Core Team, 2021).

### Study site and target-search assay

We recruited 51 volunteers (29 women and 22 men) to take part in a field-based target-detection task along a 100 m straight route marked out through a lightly shaded strip of trees on the Macquarie University campus (Sydney, Australia). The ground was covered in coarse, dry bark mulch (Fig 1B), which served as the visual background for target placement and provided a natural, visually cluttered environment with uneven texture and patchy light. We ran the experiment over five sessions between 5 and 12 May 2017, conducting trials between 11:00 and 14:00, and each participant attended one session only. In each session, we deployed 56 targets in a semi-random arrangement along both sides of the route (28 per side), comprising four replicates of each stimulus type per side (seven types in total: yellow, green, and blue stimuli in “light” and “dark” variants, plus a brown control). We placed targets at fixed lateral distances of 2, 3, 4, or 5 m to the left or right of the route centreline. We varied order and distance to avoid a predictable spatial pattern and ensured that targets of the same colour were not placed adjacent to one another. We also varied the spacing between successive targets (3 ± 0.5 m) to minimise predictability in layout. We re-randomised the order and lateral distances of targets for each session. Each participant completed a single walk along the route and thus encountered 56 targets (i.e., 56 target-presentations per participant; 2 856 target-presentations in total across all participants).

Before each trial, we provided participants with standardised instructions and showed them a single brown control target to familiarise them with the size and shape of the targets, but we did not tell them which colours they would encounter. Participants self-reported normal colour vision (i.e., no known colour-vision deficiency); we did not administer a formal colour-vision screening test, and we did not exclude any participants on this basis. Participants walked the route at a steady pace while following one of us, scanned the ground on both sides, and verbally reported the colour and side (left or right) of any targets they detected as they came adjacent to them. One of us (BR) walked just ahead of each participant to maintain pace and record detections. To characterise the light environment, at the start of each participant’s walk we recorded whether conditions were sunny, shady, or overcast, and we used a calibrated OceanView JAZ portable spectrometer with an absolute irradiance module to measure vector irradiance both outside the forest patch and at the midpoint of the route within the patch (Fig. 1e).

### Statistical analysis

To test how visual cues guide stimulus detection we used a generalised linear mixed-effect model with a binomial error family and logit link function. We specified the detection of each target in our assay as a binary response, and included target-background differences in hue, chroma, and luminance as main-effects. We also included the viewer’s distance to the target and the chromaticity of ambient light as main effects of secondary interest, to control for variation in detectability owing to the apparent size of targets and changes in ambient lighting, respectively. We specified participant identity as a random effect, to account for non-independence between records within a single trial. We also scaled and centred predictors for ease of interpretation and comparison. This singular model structure is informed by the exploratory analyses presented in White et al. (2017), the results of which underlie the explanatory design of the present work. We visually affirmed GLMM assumptions using the R package ‘DHARMa’ (Hartig 2022), and all statistical analyses were conducted using the ‘lme4’ (Bates et al. 2015) package for R v 4.3.1 (R Core Team 2021).

### Ethical note

All research was carried out under the aegis of Macquarie University’s Human Research Ethics Sub-Committee (approval #5201700333).

### Data and code availability

All data and code underlying our analyses are available via GitHub (https://github.com/invertBEACON/ms_human_detect), and will be persistently archived upon acceptance.

## Results

Across all trials, participants detected just over a third of targets overall (36% ± 1 SE), though detection rates varied sharply by stimulus type. Our background-matching controls were rarely seen (4 ± 1), as intended, while chromatic targets were detected far more often, typically close to half of presentations. Blue and green light stimuli were detected at intermediate rates (41% ± 2 and 35% ± 2, respectively), with their darker variants proving even more conspicuous (48% ± 3 and 45% ± 3). Yellow was the least visible of the chromatic set (29% ± 2), although its darker counterpart was among the most detectable (48% ± 3).

With respect to our focal hypothesis, we identified clear differences among the visual cues used to guide the detection of targets (Table 1 for full statistical results). Hue and saturation contrasts were the only visual cues that reliably predicted detection (Table 1, Fig. 2). Targets with stronger chromatic contrast against the background were detected more often, with hue contributing slightly more than saturation, although the two effects did not differ statistically (Table S1). In contrast, luminance contrast and ambient light contributed little explanatory power relative to chromatic contrasts, with weak negative coefficients suggesting only a negligible tendency for lower detection under brighter conditions. Detection declined with increasing viewing distance, confirming the expected effect of target size.

**Table 1:**
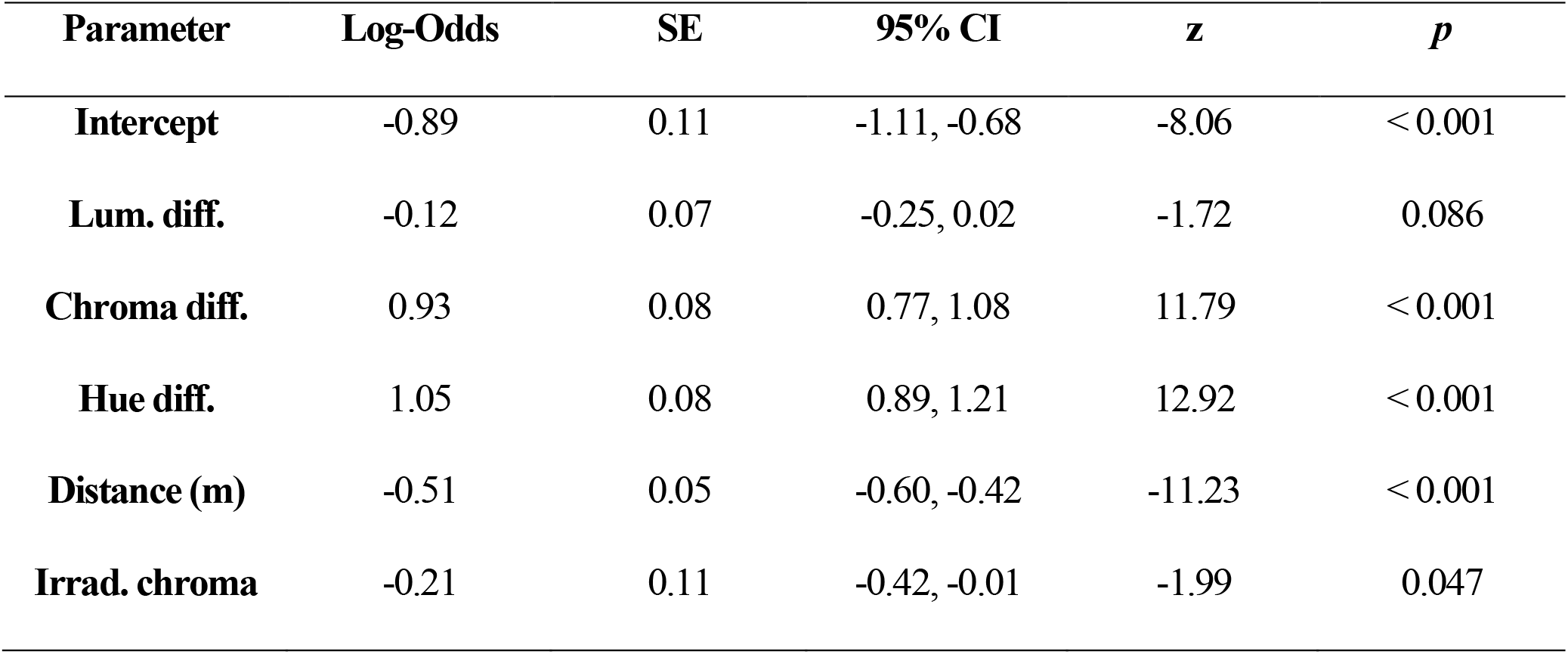
Summary of fixed effects from the generalised linear mixed-effect model predicting target detection. Log-odds estimates, standard errors (SE), 95% confidence intervals (CI), z-values, and p-values are shown for each predictor. The model included hue, chroma, and luminance contrasts (Δhue, Δchroma, Δluminance), viewing distance, and the chromaticity of ambient light as fixed effects, with participant ID as a random intercept (SD = 0.69).

| Parameter | Log-Odds | SE | 95% CI | *z* | *p* |
| --- | --- | --- | --- | --- | --- |
| **Intercept** | -0.89 | 0.11 | -1.11, -0.68 | -8.06 | < 0.001 |
| **Lum. diff.** | -0.12 | 0.07 | -0.25, 0.02 | -1.72 | 0.086 |
| **Chroma diff.** | 0.93 | 0.08 | 0.77, 1.08 | 11.79 | < 0.001 |
| **Hue diff.** | 1.05 | 0.08 | 0.89, 1.21 | 12.92 | < 0.001 |
| **Distance (m)** | -0.51 | 0.05 | -0.60, -0.42 | -11.23 | < 0.001 |
| **Irrad. chroma** | -0.21 | 0.11 | -0.42, -0.01 | -1.99 | 0.047 |

**Figure 2:**
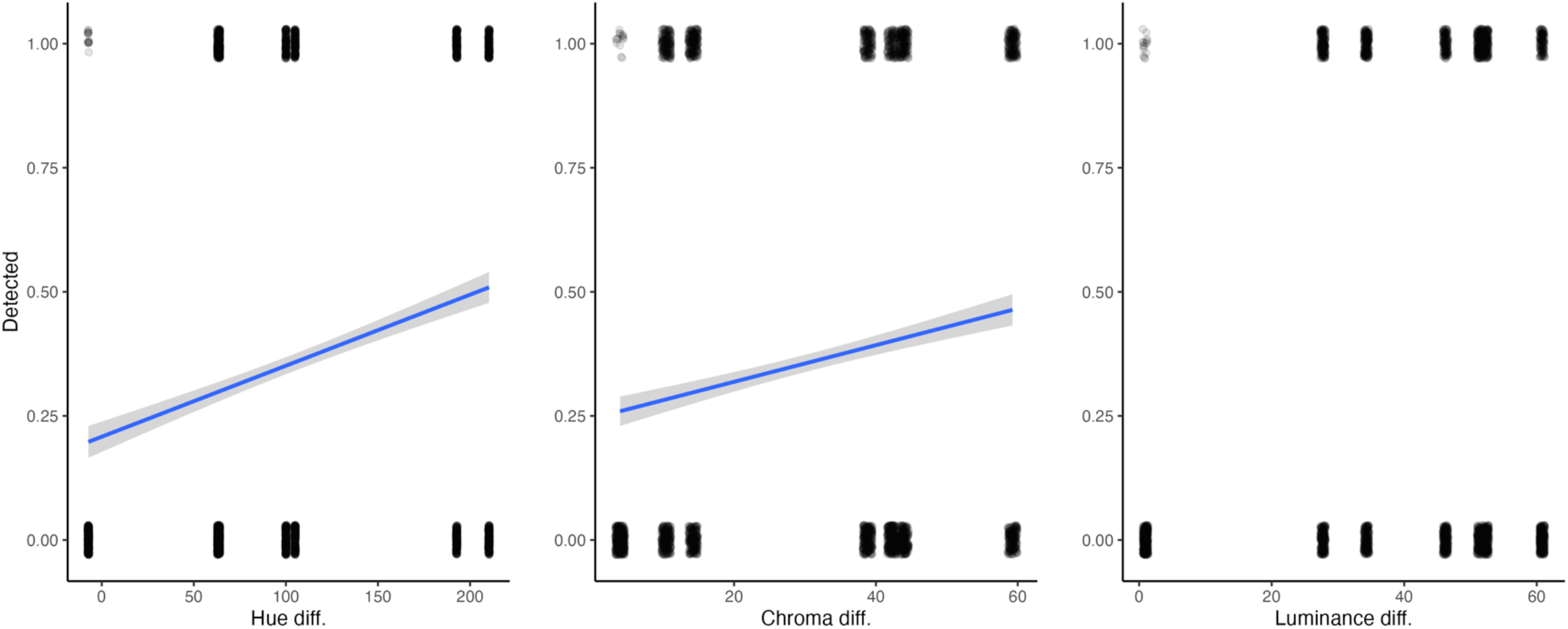
Model-predicted relationships between visual contrasts and detection probability in a test of the visual cues guiding human target detection. Lines show fitted values (± s.e.) from the generalised linear mixed model across the main effects of hue, chroma, and luminance contrasts. Each panel depicts the change in detection probability as the contrast between target and background increases along the given dimension. Hue and saturation contrasts strongly increased detections, while luminance contrast showed no reliable effect.

**Figure 3.**
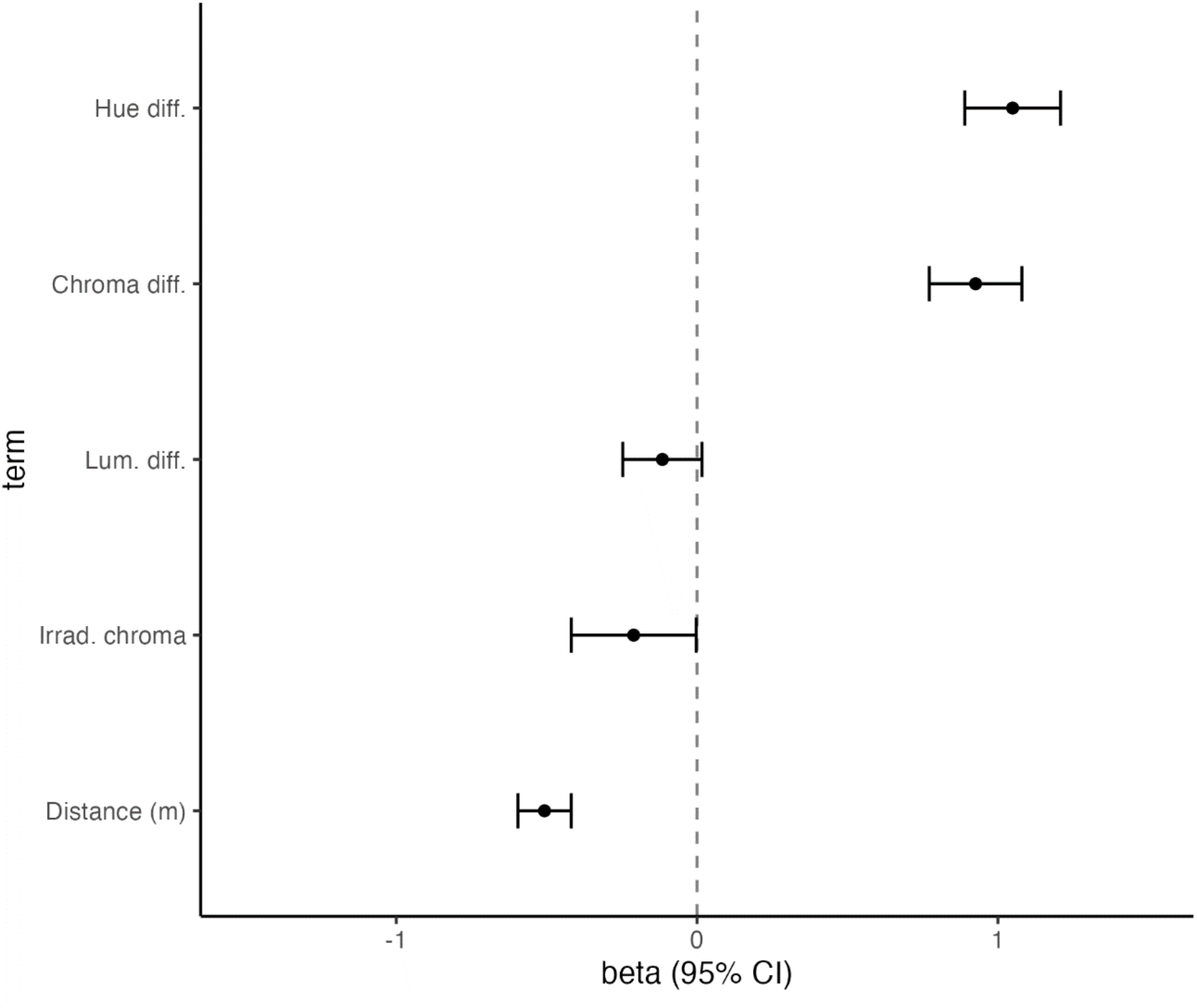
Relative importance of visual and environmental predictors in a test of the visual cues guiding human target detection. Standardised beta coefficients (β ± 95% CI) from the generalised linear mixed-effect model are shown, representing the influence of each predictor after scaling and thus allowing direct comparison of effect magnitudes. Hue and chroma contrasts emerged as the strongest positive predictors of detection, whereas luminance contrast and irradiance chroma showed weak, marginally negative effects. Detection probability also declined reliably with increasing viewing distance.

## Discussion

Natural scenes are alive with dynamic variation in colour and luminance, which presents a basic challenge for visually separating signal from noise. Here we sought to translate and extend the wealth of valuable laboratory-based insight on human visual performance to a more natural context, in a test of how key first-order visual features — hue, saturation, and luminance contrast — guide the detection and identification of stimuli in a forest-like environment. Consistent with predictions, we found that chromatic cues, specifically hue and saturation contrast, were paramount for target detection (Fig. 2). These findings broadly accord with theory and empirical findings developed in more controlled settings (Delorme et al., 2000; Kelber & Osorio, 2010), though also speak to a complex picture of differential cue-use depending on the visual task and context.

Surprisingly, luminance contrast contributed little explanatory power relative to chromatic contrasts. The primacy of chromatic contrasts in guiding target identification in our study is consistent with the stability of colour-based cues in natural scenes (Kelber & Osorio, 2010; Vorobyev, 2004). Chromatic edges can be as informative as luminance edges in natural scenes and contribute to object-contour perception (Hansen & Gegenfurtner, 2009, 2017), and colour information supports the rapid recognition of natural scenes (Gegenfurtner & Rieger, 2000). Whereas the luminance of objects can vary by orders of magnitude with passing clouds or the dappling of light through canopies, the hue and, in particular, saturation of stimuli is a comparatively stable guide to their character (Osorio & Vorobyev, 2005; Vorobyev, 2004). The visual backgrounds in our study were naturally ‘noisy’, especially in lightness. While the desaturated browns that comprise the bark background present a palette which is relatively constrained in both hue and saturation, the spatially and temporally changeable intensity of veiling light through the canopy will naturally inhibit the identification of stimuli on the basis of luminance alone (Endler, 1993). When considered in concert with higher-level processes like colour constancy and perceptual normalisation for stabilising colour appearance amidst noise (Gegenfurtner & Kiper, 2003; Geisler, 2008), and humans’ known ability to flexibly attend to salient cues (Frey et al., 2007), the predictive value of chromatic contrasts in our assays stands to reason.

This central finding affirms the results of laboratory-based efforts, though it also highlights the richer context-dependence of cue use in natural settings. In standard image-based tasks, humans appear to use first-order cues in a highly flexible manner, depending on scene structure and task demands (Frey et al., 2007, 2008; Parkhurst et al., 2002). In naturalistic images such as forests, colour information often dominates luminance in guiding overt attention, as estimated by eye fixations (Frey et al., 2008). In contrast, artificial scenes — such as urban environments or fractal patterns — may favour luminance cues due to their pronounced edges and strong spatial gradients (Parkhurst et al., 2002). Yet even within these categories, contributions of colour can be significant (Parkhurst et al., 2002), and in some cases, first-order cues altogether fail to predict visual attention (Frey et al., 2007). These inconsistencies likely stem from subtle differences in image sets, task instructions, and model parameterisation, underscoring both the complexity of cue use and the limits of strictly ‘bottom-up’ approaches to understanding the ecology of human colour vision.

Our knowledge of realised, rather than potential, visual performance ultimately demands experiments conducted in naturalistic conditions. Direct tests in humans, however, are few. White et al. (2017) describes one such recent attempt, though it leveraged a pre-existing dataset using humans as model predators of colour-polymorphic frogs. In only partial agreement with the present study, our results in that instance ranked hue and luminance contrasts as primary, with saturation being non-predictive of detections. A key distinction, and likely explanation, is the difference in stimuli: White et al. (2017) was constrained to a naturalistic colour palette in keeping with the frogs’ ecology (Rojas et al., 2014), limiting the range of chromatic variation. Our stimuli, in contrast, were purpose-designed to span a broader range of hue, saturation, and luminance values, affording greater variation in the contrasts available to viewers (Fig. 1d). Since humans can flexibly attend to the most diagnostic cues for a given task (Oliva & Schyns, 2000), the primacy of luminance in the earlier study may simply reflect a context in which saturation was less informative — rather than any inherent inferiority of that cue.

Naturalistic tests of visual performance are, understandably, far more common in non-humans, and our findings thus represent an early anchor point for humans within this rich corpus (Kelber & Osorio, 2010; Vorobyev, 2004). In New World monkeys, for example, chromatic cues are important in guiding attention and identifying salient stimuli, though fast achromatic channels continue to dominate motion processing and broad scene analysis (Gegenfurtner & Kiper, 2003; Hubel & Livingstone, 1987). Studies in other primates echo this balance, suggesting a conserved visual architecture in which luminance is fast and coarse, while colour is stable and informative (Vorobyev, 2004). In that sense, our findings support a view of shared perceptual foundations across primates, with chromatic cues — especially hue and saturation — acting as robust guides in visually complex environments. While our focus was limited to first-order contrasts, humans possess a richer perceptual toolkit, incorporating higher-order processes such as pattern recognition, figure-ground segregation, and selective attention (Frey et al., 2007; Párraga et al., 2000). These lend additional flexibility and nuance to cue use, allowing humans to differentially weight chromatic and achromatic channels depending on the task at hand. This layered perceptual capacity — grounded in shared mammalian architecture yet elaborated in humans — invites comparative work exploring how different taxa resolve signal from noise in natural environments, and how such strategies have evolved in relation to ecological niche, sensory constraints, and behavioural flexibility.

Human colour vision remains both deeply familiar and deeply complex; a system whose basic physiology is well characterised, yet whose real-world performance remains underexplored. Our study helps bridge this gap, offering rare field-based insight into how first-order chromatic and achromatic contrasts guide detection under natural conditions. That chromatic contrasts emerged as primary here reflects not only their stability in noisy environments, but also the contextual flexibility of human perception. Indeed, a key lesson is that cue use is not fixed: it varies with the demands of the task, the structure of the background, and the characteristics of the stimulus. This has implications for how we design signals, interpret camouflage, and model other animals’ perceptual experiences. There remains substantial need — and opportunity — for further work in this vein, as we continue the essential task of translating lab-based insight into the ecologically grounded realities of vision in the wild.

## Acknowledgements

TEW was supported by an Australian Research Council DECRA fellowship (DE230100087). DJK was supported by the Human Frontier Science Program (RGP023/2023). BR was supported by a mobility grant from the Research Council of the University of Jyväskylä. Both BR and JM were funded by the Finnish Centre of Excellence in Biological Interactions (Research Council of Finland, project no. 284666 to JM).

