## supplementary material for "Chromatic contrasts predict human target detection in a natural search task"

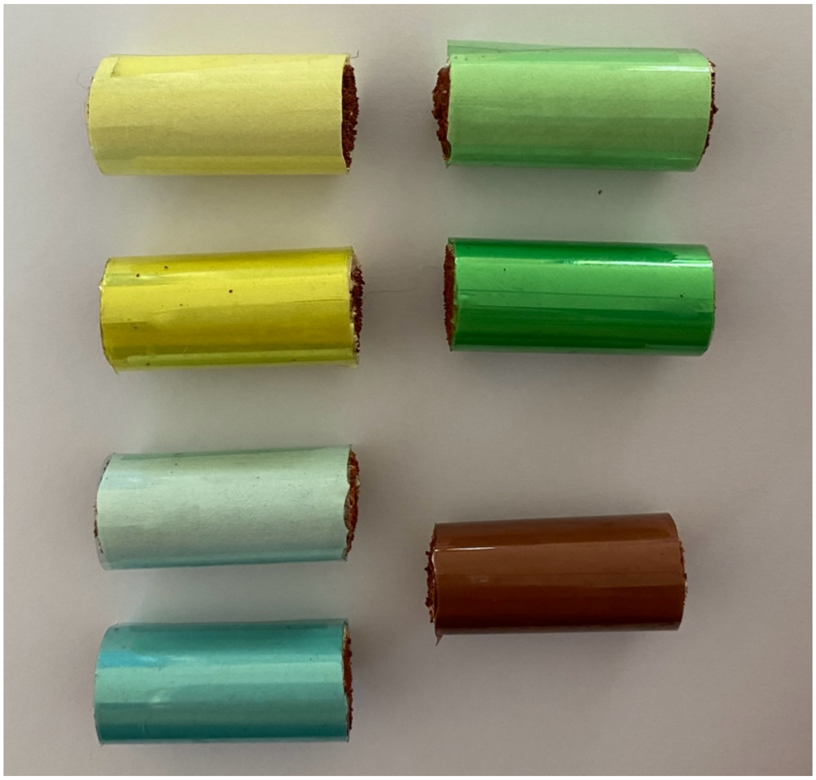

**Figure S1:** Photographs of the experimental targets used in a test of the visual cues guiding human target detection, corresponding to the spectral representations shown in Figure 1. The seven stimuli comprised three light (single-layer) and three dark (double-layer) chromatically matched targets (yellow, green, blue), along with a background-matching control (brown). The contrast in lightness between the "light" and "dark" variants is evident here, illustrating the range of chromatic and achromatic cues presented to participants.

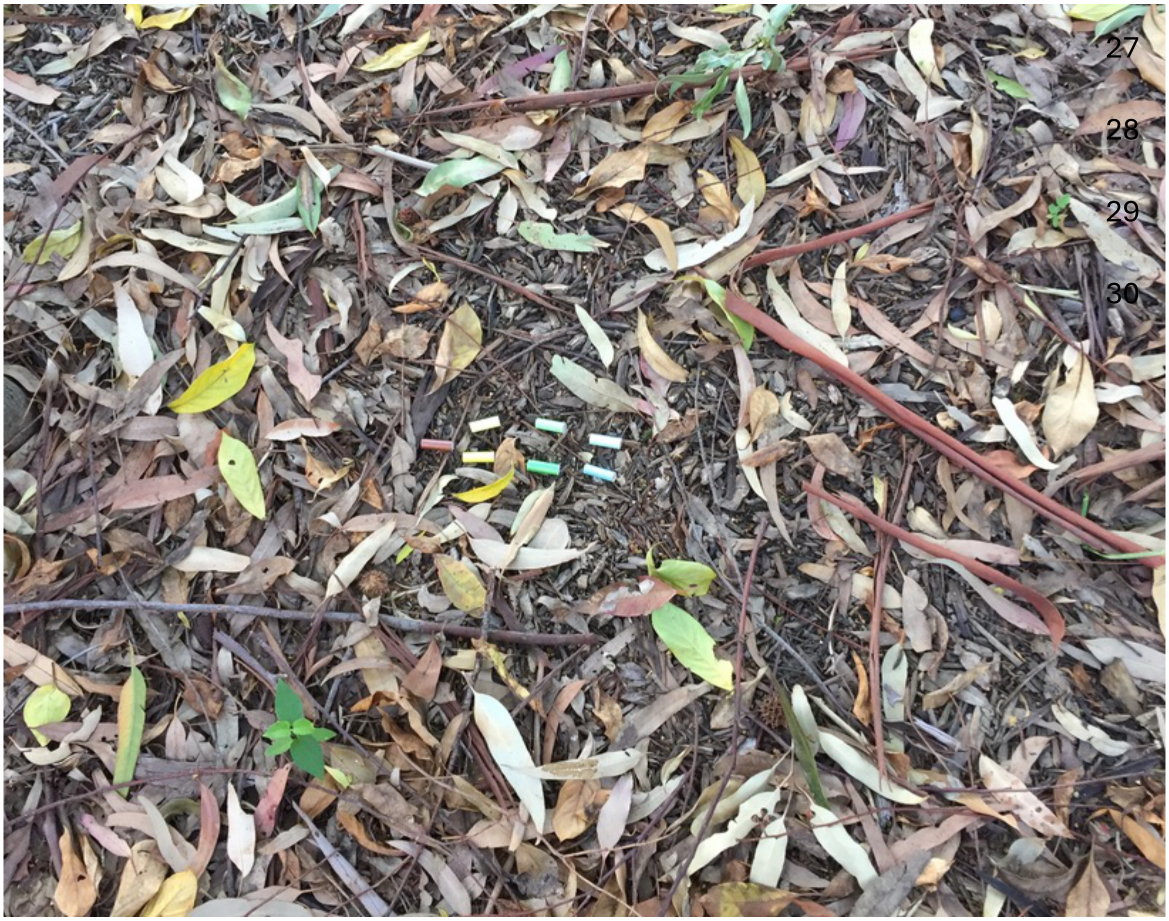

**Figure S2:** Heuristic example of the natural search background and target appearance *in situ* during field detection trials. Coloured dowel targets (centre) are shown resting on the substrate surface, illustrating the visual clutter and patchy texture against which participants searched for targets and reported detections. Targets comprised chromatically matched yellow, green, and blue stimuli presented as “light” (single-layer) and “dark” (double-layer) variants, plus a background-matching brown control.
